# The ELT-3 GATA Factor Specifies Endoderm in *Caenorhabditis angaria* in an ancestral gene network

**DOI:** 10.1101/2022.05.25.493523

**Authors:** Gina Broitman-Maduro, Morris F. Maduro

## Abstract

Endoderm specification in the nematode, *C. elegans*, occurs through a well-characterized pathway that is initiated by maternally provided SKN-1/Nrf, and with additional input from POP-1/TCF, which activates the GATA factor cascade MED-1,2 → END-1,3 → ELT-2,7. Orthologues of the MED and END factors, and ELT-7, are found only among nematodes of the Elegans Supergroup consisting of species closely related to *C. elegans*, which raises the question of how gut is specified in their absence. In this work, we investigate gut specification outside the Elegans Supergroup. We find that the *C. angaria* and *C. portoensis* orthologues of the *elt-3* GATA factor gene are expressed in the early E lineage, just before their *elt-2* orthologues. In *C. angaria*, both *Can-pop-1(RNAi)* and *Can-elt-3(RNAi)* result in a penetrant ‘gutless’ phenotype. *Can-pop-1* is necessary for *Can-elt-3* activation, showing that it acts upstream. When introduced into *C. elegans* as transgenes, overexpressed *Can-elt-3* is sufficient to specify gut, while *Can-elt-2* can rescue gut differentiation under the control of its own promoter. Our results demonstrate an ancestral mechanism for gut specification and differentiation in *Caenorhabditis* involving a simplified gene network consisting of POP-1 → ELT-3 → ELT-2.

**Summary statement:** Specification of the gut progenitor E in a distant relative of *C. elegans* uses a different GATA factor, ELT-3, suggesting that the ancestral network was simpler.

## Introduction

Gene regulatory networks drive cell specification events in metazoan systems, and are subject to large-and small-scale changes over evolutionary time (True and Haag, 2001; Davidson and Levine, 2008). One of the most studied networks in model systems is that which specifies the *C. elegans* gut cell progenitor E and promotes intestine differentiation in its descendants (Fig. 1A,B). At the top of the network, the maternal SKN-1/b-ZIP factor, acting partially through its zygotic effectors MED-1,2, along with the maternal Wnt/β-catenin asymmetry pathway through its effector POP-1/TCF, activate early E lineage expression of the *end-3* and *end-1* genes (Bowerman et al., 1992; Lin et al., 1995; Rocheleau et al., 1997; Thorpe et al., 1997; Maduro et al., 2001; Maduro et al., 2002; Shetty et al., 2005; Bhambhani et al., 2014). Downstream of these transiently expressed factors, *elt-2* and its paralogue *elt-7* drive gut development and differentiation and expression is maintained through adulthood (Fukushige et al., 1998; Sommermann et al., 2010; Dineen et al., 2018).

**Fig. 1.**
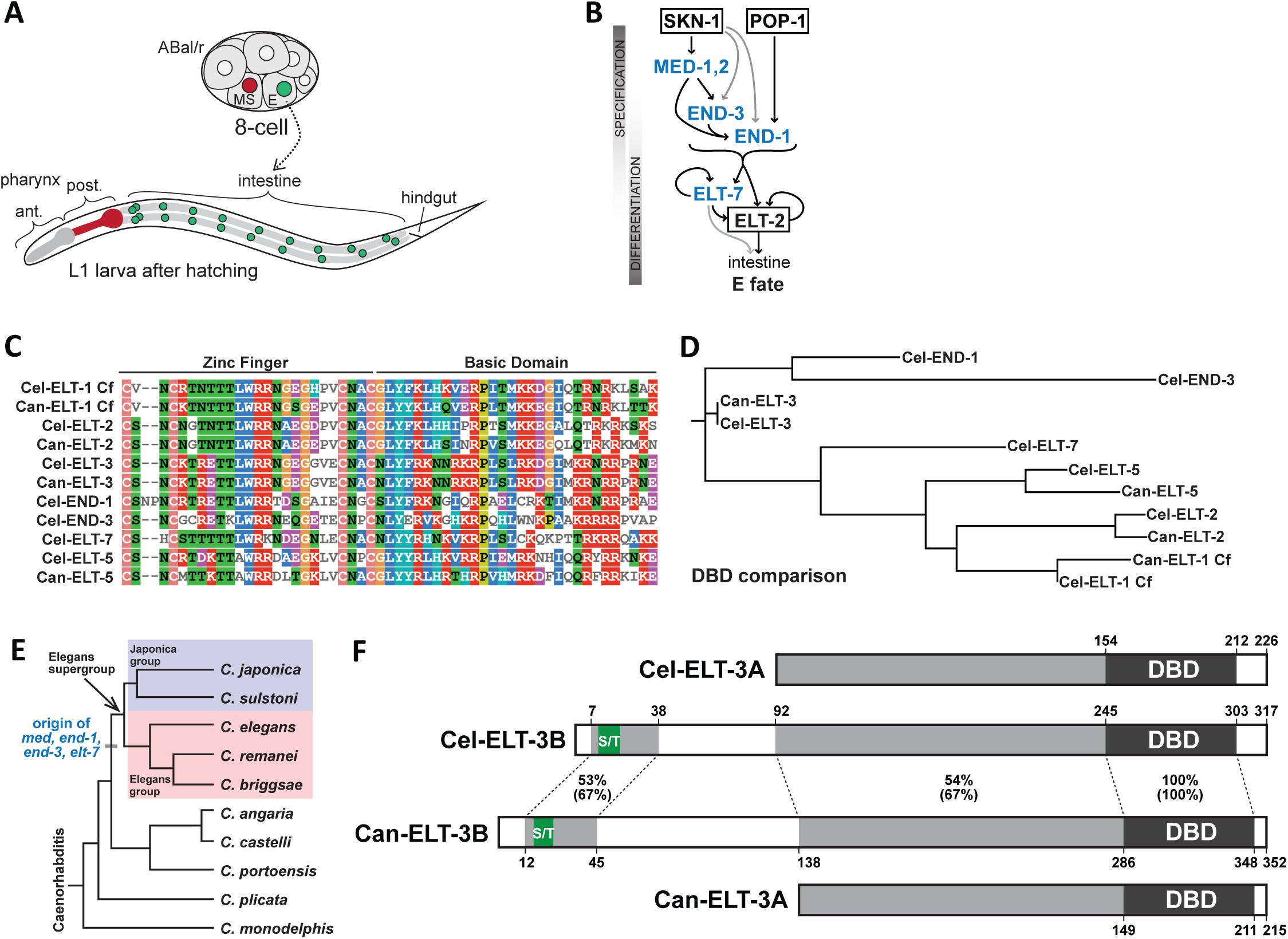
Background information on the E blastomere, the endoderm gene network, and GATA factor conservation. (A) Lineal origin of the gut precursor E in *Caenorhabditis*. E (nucleus shaded green) is located posteriorly and ventrally at the 8-cell stage and gives rise to 20 descendants that form the juvenile intestine (nuclei shaded green). The sister cell of E, called MS (red nucleus), makes mesodermal cell types that include the posterior cells of the pharynx, also shaded red (Sulston et al., 1983). (B) Simplified diagram of the endoderm specification network from *C. elegans* (Maduro, 2017). The factors that are absent in more distant relatives of *C. elegans* are shaded in blue. (C) Alignment of the DNA-binding domains (C4 Zinc Finger and Basic Domain) for the canonical GATA factors in *C. elegans* and *C. angaria*. The colored blocks were generated by MView Multiple Sequence Alignment (https://www.ebi.ac.uk/Tools/msa/mview/). (D) RAxML-NG tree of the DNA-binding domains shown in (C) generated using CIPRES Gateway (https://www.phylo.org/) similar to trees made in a prior work (Eurmsirilerd and Maduro, 2020). The *C. elegans* genome contains two additional GATA factor paralogs, *elt-4*, which is a partial duplication of *elt-2* that lacks a detectable function, and *elt-6*, a paralogue of *elt-5/egl-18* (Koh and Rothman, 2001; Fukushige et al., 2003). (E) The *med, end*, and *elt-7* factors likely originated at the base of the Elegans supergroup, a sub-clade of the *Caenorhabditis* genus that includes the Japonica and Elegans groups (Maduro, 2020). (F) Alignment of *C. elegans Cel-ELT-3B* long and short *Cel-ELT-3A* protein isoforms with the predicted orthologues from *C. angaria*, with results of protein-protein BLAST comparison. The DNA-binding domains are identical, while the immediate upstream region common to both short isoforms shows 54% identity (67% similarity). A short 21-23 amino acid region is conserved at the amino end that includes a poly-serine/threonine region (S/T). In *C. elegans* the Poly-S/T region has sequence SSTSSSDS (7/8 amino acids are serine or threonine, 6/8 are serine), and in *C. angaria*, it is SPHSSTDTSS (7/10 amino acids are serine or threonine, 5/10 are serine). These sequences occur in other endodermal GATA factors, however their significance is unknown (Eurmsirilerd and Maduro, 2020; Maduro, 2020).

As would be expected for a network with so many genes, the factors between SKN-1 and *elt-2* demonstrate complex patterns of partial or complete redundancy. Some genes, like *end-1* and *elt-7*, can be individually deleted with no apparent phenotype, while other combinations result in large fractions of embryos lacking gut (Maduro et al., 2005a; Sommermann et al., 2010). The most penetrant zygotic defect in gut results from mutants lacking both *end-1* and *end-3* together, which fail to specify gut 100% of the time (Zhu et al., 1997; Maduro et al., 2005a; Owraghi et al., 2010). Generally, any genotype leading to partially compromised specification leads to a loss of robustness of *elt-2* activation and a failure to develop a completely normal intestine in terms of both gut cell number and metabolic function (Maduro et al., 2007; Raj et al., 2010; Maduro et al., 2015).

The zygotic endoderm genes all encode GATA factors, a family of transcription factors that bind the canonical sequence HGATAR (Lowry and Atchley, 2000; Wiesenfahrt et al., 2015; Du et al., 2016). The MED factors are an unusual divergent subfamily, binding to a related RAGTATAC core sequence (Broitman-Maduro et al., 2005; Lowry et al., 2009). The canonical *C. elegans* GATA factors, including the endodermal END-1,3 and ELT-2,7 factors, have similar DNA binding domains (DBDs) (Fig. 1C,D). In two interesting demonstrations of functional overlap, forced early endoderm expression of ELT-2, using the *end-1* promoter (*end-1p*::ELT-2), can functionally replace the upstream function of *end-3, end-1*, and *elt-7*; and the function of all of *end-1,3* and *elt-2,7* can be replaced by a double transgenic combination of *end-1p*::ELT-7 and *elt-2p*::ELT-7 (Wiesenfahrt et al., 2015; Dineen et al., 2018), suggesting that there is a high degree of functional overlap among GATA factors. The overlap does have limits, however, as high copy numbers of these transgenes are required for their function, suggesting there are factor-specific functions that depend on regions upstream of the DBDs (Wiesenfahrt et al., 2015; Dineen et al., 2018).

As essential as the endoderm network is for *C. elegans*, there are no orthologues of the *med, end*, and *elt-7* genes outside of the Elegans Supergroup of species in *Caenorhabditis*, suggesting they evolved over a short time period at its base (Maduro, 2020) (Fig. 1E). Most nematodes in the broad clade of Rhabditids that includes the genus *Caenorhabditis* have only four ‘core’ GATA factors that are orthologous to factors found in *C. elegans* (Eurmsirilerd and Maduro, 2020). Aside from ELT-2, there are the ELT-1 and ELT-3 factors, which both function in hypodermal specification, and ELT-5/EGL-18, which specifies hypodermal cells in the lateral seam (Page et al., 1997; Gilleard and McGhee, 2001; Koh and Rothman, 2001; Koh et al., 2002).

In this work, we examine gut specification outside of the Elegans Supergroup using *C. angaria* (Felix et al., 2014). This species has several advantages for study, including its robust growth under laboratory conditions like those for *C. elegans*, and the fact that it is has been used in comparative studies by multiple laboratories, with some examples here (Jud et al., 2007; Kuntz et al., 2008; Brauchle et al., 2009; Nuez and Felix, 2012; Barkoulas et al., 2016; Macchietto et al., 2017). RNAi has shown some success in *C. angaria* (Nuez and Felix, 2012). Embryos of *C. angaria* resemble those of *C. elegans* and undergo a similar development in just over 11 hours at 24°C with minor variations in the times at which particular milestones are reached (Macchietto et al., 2017). Using a combination of *C. elegans* transgenics, single-molecule FISH detection, and RNA interference in *C. angaria*, we present multiple lines of evidence that gut specification in *C. angaria* occurs by the *Can-POP-1-*dependent activation of *Can-elt-3*. In turn, *Can-ELT-3* activates *Can-elt-2* to drive gut differentiation. The results suggest an evolutionary origin of the endoderm gene regulatory network in the Elegans Supergroup from a simpler ancestral GATA factor cascade.

## Results

### Testing requirements for maternal Can-SKN-1 and Can-POP-1

We began by testing the possibility that the *C. angaria* orthologues of SKN-1 and POP-1 might play a role in gut specification. The *Can-skn-1* and *Can-pop-1* orthologues appear to be maternally expressed, as single-embryo RNA-Seq experiments in zygotes and 2-cell stage embryos recovered transcripts for these (Macchietto et al., 2017). To deplete function of these genes individually, we tried RNA interference using both feeding-based RNAi and direct gonadal injection of dsRNA. Using both approaches, we observed a penetrant embryonic lethality with *Can-pop-1(RNAi)*. After at least 24h of growth of L4/adult animals on *Can-pop-1* dsRNA-expressing bacteria, 90% of embryos (n=252) showed a uniform arrest phenotype, at onefold elongation with several hundred nuclei but no morphogenesis (Fig. 2C). RNAi by dsRNA injection resulted in the same phenotype, though only 149/234 (64%) of progeny embryos were affected, likely because we injected only a single gonad arm per female to favor survival. All arrested *Can-pop-1(RNAi)* embryos lacked differentiated gut, as evidenced by the lack of birefringent gut granules. This phenotype was immediately striking to us, as RNAi of *pop-1* in *C. briggsae* resulted in a very similar phenotype (Lin et al., 2009). Based on the similarity of phenotype to *C. briggsae*, therefore, we hypothesized that *Can-pop-1* is required for gut to be specified in *C. angaria*, acting upstream of specification as it does in *C. briggsae*. (We will directly test this hypothesis later in this work.)

**Fig. 2.**
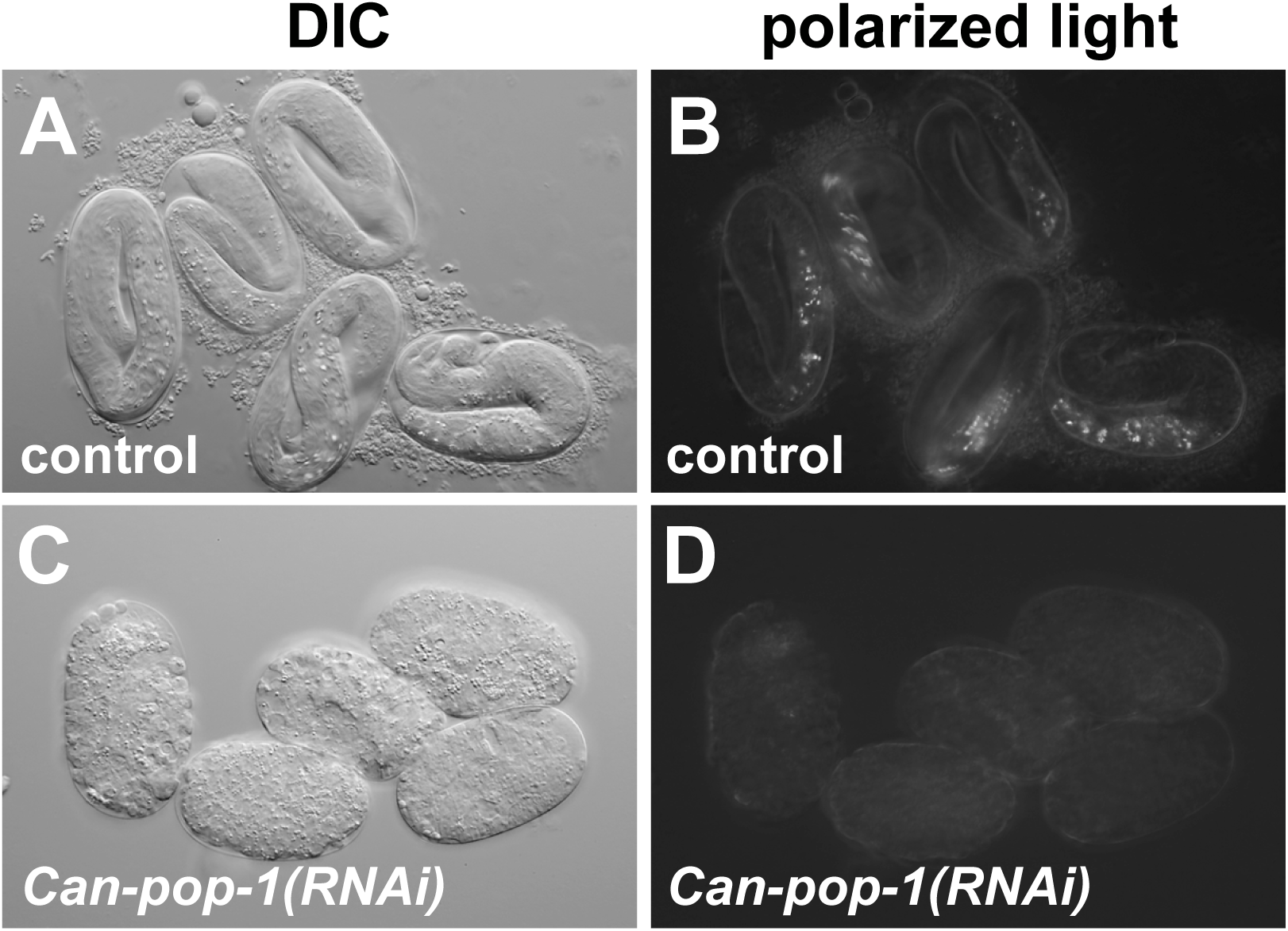
RNA interference of *Can-pop-1*. (A) Control untreated RGD1 *C. angaria* embryos shortly before hatching, by differential interference contrast (DIC). (B) Birefringent gut granules in controls visualized by polarized light. 98/102 (96%) control embryos elongated to threefold stage and 100% contained gut granules. (C) DIC image of onefold arrested embryos of *Can-pop-1(RNAi)*. 226/252 (90%) progeny embryos arrested following RNAi by feeding treatment. Arrested embryos lacked differentiated intestine and arrested without significant morphogenesis. (D) Interfered *Can-pop-1(RNAi)* embryos lack gut granules. A *C. angaria* embryo is approximately 50 μm long. Anterior is to the left, and dorsal is up in most cases in this and other figures.

We looked for evidence of a fate transformation from E in *Can-pop-1(RNAi)* embryos. In our *C. briggsae* studies, *Cbri-pop-1(RNAi)* resulted in the E cell adopting the fate of its sister, MS (Lin et al., 2009). We tested for such a transformation by looking for ectopic pharyngeal tissue by staining for *Can-myo-2* by smiFISH, similar to the *in situ* hybridization approach used in our *C. briggsae* study (Okkema et al., 1993; Lin et al., 2009). However, we did not observe any evidence of extra *Can-myo-2* expression (data not shown).

We next attempted to knock down *Can-skn-1* using RNAi by feeding and RNAi by injection as for *Can-pop-1*. Whereas we easily observed a highly penetrant embryonic arrest with *Can-pop-1(RNAi)*, RNAi targeting *Can-skn-1* resulted in no apparent phenotype (n>200), using both dsRNA injection and RNAi by feeding and with two different targeting sequences. Occasionally, unusual embryos or larvae were observed in less than 5% of progeny that had various morphological or elongation defects, however these were also observed at a similar frequency following control dsRNA injection, control RNAi by feeding, or no treatment. Because *C. angaria* is a male-female species, these rare embryos likely result from a reduction in developmental robustness due to inbreeding depression (Nuez and Felix, 2012).

Regardless, even these rare abnormal animals all contained differentiated intestine as visualized by gut granule birefringence. Hence, we were unable to detect an obvious phenotype for *Can-skn-1(RNAi)* in development, let alone in gut specification, in stark contrast to the essential requirement for *Cel-skn-1* in *C. elegans*. (A control for knockdown of *Can-skn-1* will be described later.)

### Expression of Can-elt-2 as a transgene in C. elegans

Because orthologues of the intermediate endodermal GATA factors from the Elegans Supergroup are absent in *C. angaria*, we next examined *Can-elt-2*. ELT-2 is widely conserved among nematodes (Eurmsirilerd and Maduro, 2020). The parasite *H. contortus*, a parasitic nematode within the Rhabditina, encodes an apparent *elt-2* orthologue that can promote gut fate when overexpressed in *C. elegans* (Couthier et al., 2004). We therefore expected that the putative *Can-elt-2* gene functions in differentiation of the *C. angaria* gut. We hypothesized that if the entire *Can-elt-2* gene was introduced into *C. elegans*, it may be capable of intestinal expression, driven (at least) by *C. elegans* ELT-2 and ELT-7 acting through autoregulatory GATA sites in the *Can-elt-2* promoter (Fukushige et al., 1999; Wiesenfahrt et al., 2015).

We amplified the *Can-elt-2* gene with 5.0 kbp of its upstream flanking DNA, the entire coding region including introns, and 231 bp downstream of the stop codon. We inserted the coding region for GFP just before the stop codon. In a wild-type background, the *Can-ELT-2::GFP* transgene is indeed expressed in intestinal nuclei in *C. elegans*, starting in the early embryo and continuing through adulthood (Fig. 3A,B,D,E), similar to expression of a *Cel-elt-2* reporter (Fig. 3C). Curiously, we regularly observed a small subnuclear spot of *Can-ELT-2::GFP* in nuclei, which was particularly prominent in the gut of young adults (Fig. 3D; 36% of 547 gut nuclei examined in 20 worms). These were reminiscent of spots observed from autoregulatory interaction of *C. elegans* ELT-2::GFP protein with the *elt-2* promoter DNA on a multicopy transgene array (Fukushige et al., 1999). The nuclear spots suggest, therefore, that the *C. angaria elt-2* gene is capable of positive autoregulation, similar to *Cel-elt-2* (Fukushige et al., 1999). Consistent with its expected intestine-specific role, we did not observe expression of *Can-ELT-2::GFP* outside of the E lineage.

**Fig. 3.**
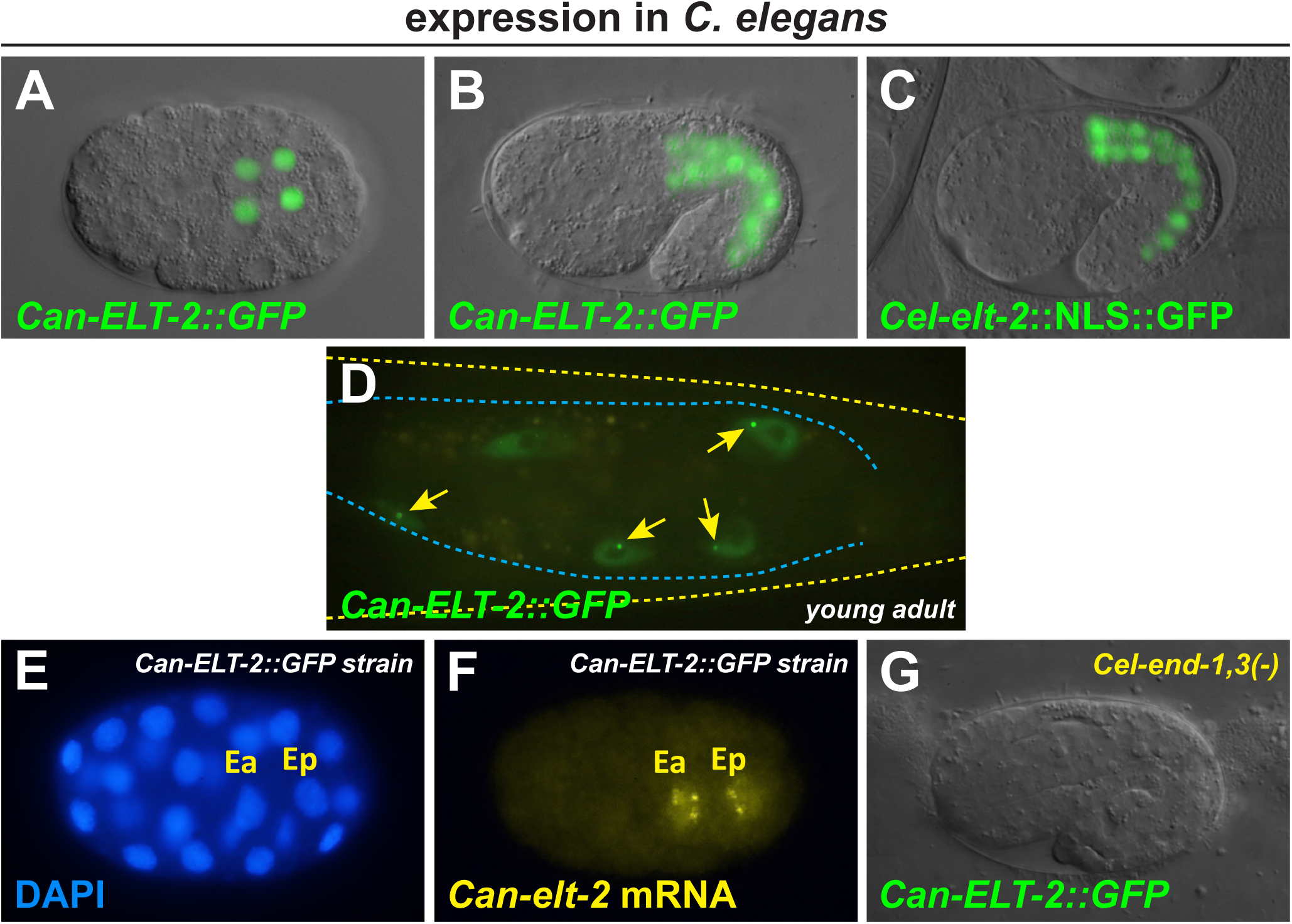
Expression of *Can-ELT-2::GFP* in *C. elegans* under the control of its own promoter. (A) Detection of *Can-ELT-2::GFP* at the 4E stage. (B) Expression in a 1.5-fold stage embryo. (C) Expression of a *Cel-elt-2*::NLS::GFP reporter transgene in an embryo slightly younger than that shown in (B). In this image the expression was much brighter so a much shorter exposure was used to image the GFP. (F,G) smiFISH probes detect nascent *Can*-*ELT-2::GFP* mRNA at the late 2E stage. (H) *Can-ELT-2::GFP* expression fails to occur in the absence of *end-1* and *end-3*. In panels A,B,C and G, a DIC image was overlaid with a fluorescence image. A *C. elegans* embryo is approximately 50 μm long.

### Can-ELT-2::GFP rescues a C. elegans elt-2; elt-7 double mutant

The early activation of *Can-ELT-2::GFP*, and its possible positive autoregulation, both suggested that the *Can-ELT-2::GFP* transgene could substitute for the endogenous *Cel-elt-2* gene. We introduced the *Can-ELT-2::GFP* transgene into a *C. elegans elt-2(ca15); elt-7(tm840)* double null mutant background, to eliminate both *Cel-elt-2* and any overlapping function contributed by *Cel-elt-7*. Double mutant *elt-2; elt-7* animals arrest as first-stage larvae with incompletely developed intestines (Sommermann et al., 2010). As anticipated, the *Can-ELT-2::GFP* transgene rescued the larval lethality of the strain to viability at high efficiency (>95% of transgenic animals). Expression of *Can-ELT-2::GFP* in the *elt-2; elt-7* strain was indistinguishable from expression in a wild-type background. The ability of *Can-ELT-2::GFP* to rescue a *Cel-elt-2; elt-7* double mutant confirms that *Can-ELT-2::GFP* can drive gut development in *C. elegans* independently of endogenous *Cel-*ELT-2 and *Cel-*ELT-7, and furthermore, that expression must be maintained once initiated.

### Expression of Can-ELT-2::GFP in C. elegans requires prior gut specification by END-1,3

The activation of *Can-ELT-2::GFP* in *C. elegans*, in the absence of endogenous *Cel-elt-2* and *Cel-elt-7*, imply that expression is activated by upstream gut specification factors. To more precisely determine when the *Can-ELT-2::GFP* transgene was being activated in *C. elegans*, we detected nascent transcripts using smiFISH (Tsanov et al., 2016; Parker et al., 2021). As shown in Fig. 3E,F, the earliest transcripts of *Can-ELT-2::GFP* were detected in the nuclei of Ea and Ep just after the E daughters have ingressed into the embryo, slightly earlier than endogenous *Cel-elt-2* transcripts become detectable, but later than when *Cel-end-3* transcripts first appear (Raj et al., 2010; Nair et al., 2013). This timing suggested that *Can-ELT-2::GFP* was being activated by END-3 and END-1. To test this hypothesis, we crossed the *Can-ELT-2::GFP* transgene into a double mutant *end-1(ok558) end-3(ok1448)* strain that is maintained by an *end-3(+)* array marked with *unc-119*::mCherry (Owraghi et al., 2010). We examined embryos in which mCherry was absent, hence are double mutant *end-1 end-3*, but which express *unc-119*::CFP, confirming the presence of the *Can-ELT-2::GFP* array. We found that *Can-ELT-2::GFP* expression was abolished in such embryos (Fig. 2H). Of 112 embryos lacking *unc-119::mCherry*, all (100%) lacked *Can-ELT-2::GFP* expression and any visible evidence of gut. To test whether END-1 by itself was sufficient to activate *Can-ELT-2::GFP*, we introduced the transgene into an *end-3(ok1448)* single mutant. In this background, *Can-ELT-2::GFP* was still expressed in transgenic embryos that made gut (data not shown). We conclude that expression of the *Can-ELT-2::GFP* transgene in *C. elegans* requires prior specification of gut by *end-1* and *end-3*, a striking result because *C. angaria* lacks orthologues of these genes.

### C. angaria elt-3 is expressed in the early E lineage

To explain the activation of *Can-ELT-2::GFP* in *C. elegans* by END-1,3, we speculated that within *C. angaria*, endogenous *Can-elt-2* might be activated by another GATA factor. Hence, we used smiFISH to examine endogenous expression of *elt-1, elt-2 elt-3*, and *elt-5* (Figs. 4,5). As predicted by expression of *Can-elt-2::GFP* in *C. elegans*, we detected *Can-elt-2* transcripts starting at the 2E stage, after the two E daughters have moved into the interior of the embryo, and continuing in the E lineage and intestine at older stages (Fig. 4M-P). This pattern is similar to what we observed for the *Can-ELT-2::GFP* transgene in *C. elegans* by smiFISH, and aside from the earlier onset, similar to expression of endogenous *Cel-elt-2* (Fig. 4Q-T). Because we did not see transcripts for *Can-elt-2* in embryos younger than the late 2E stage, this suggested to us that whatever activates *Can-elt-2* in *C. angaria* cannot be *Can-ELT-2* itself.

**Fig. 4.**
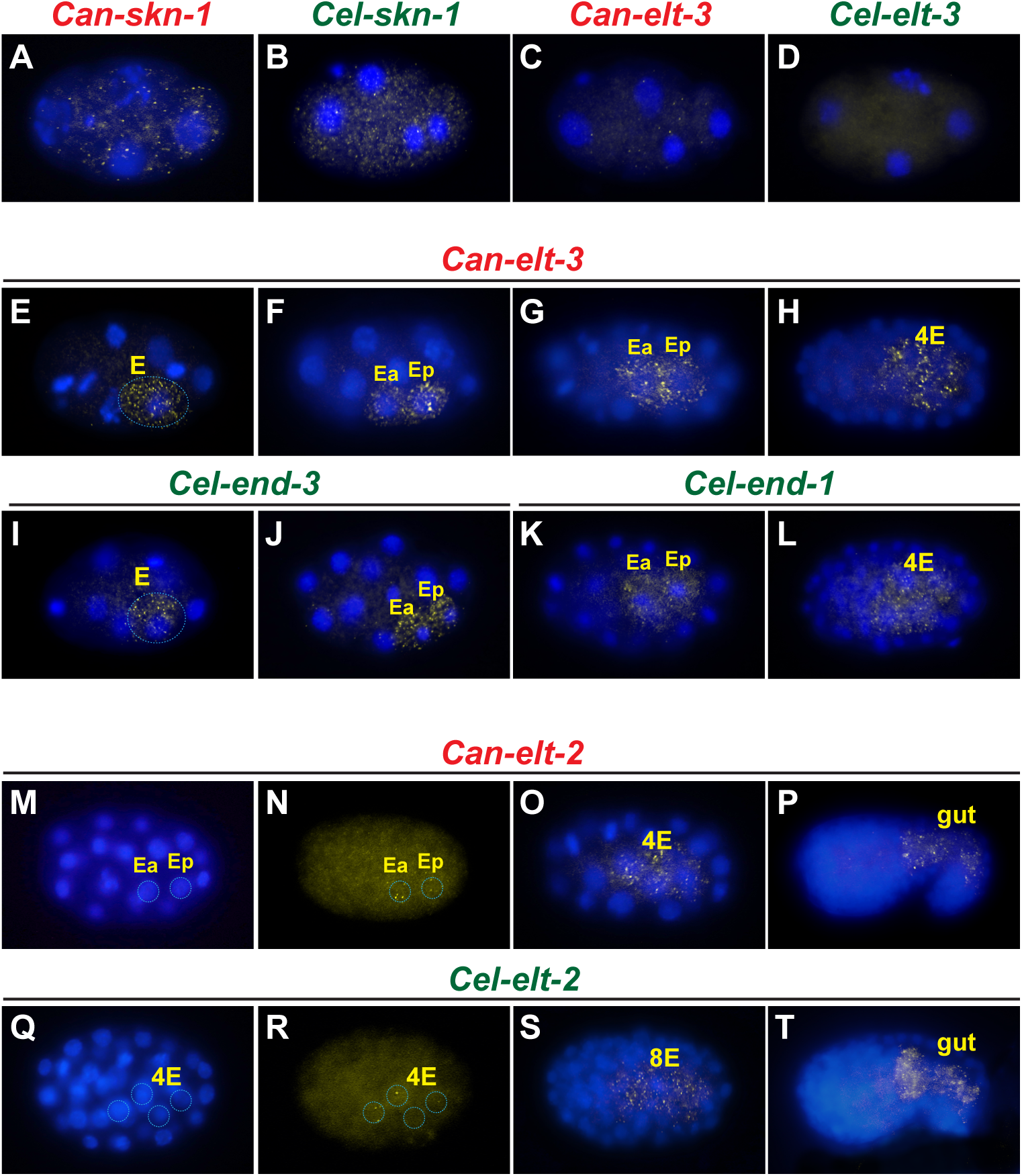
Expression of genes in *C. angaria* and *C. elegans* by smiFISH. (A) Maternal transcripts of *Can-skn-1* at the 4-cell stage. (B) Maternal *Cel-skn-1* at the 4-cell stage. (C) Apparent low-level maternal transcripts of *Can-elt-3* at 4-cell stage. (D) Lack of maternal transcripts of *Cel-elt-3* at 4-cell stage. (E-H) Expression of *Can-elt-3* in the early E lineage from E through to 4E. In (E) the E cell is outlined. (I,J) Earliest *Cel-end-3* expression from E to 2E. In I, the E cell is outlined. (K,L) Earliest *Cel-end-1* expression from 2E to 4E. (M,N) Onset of *Can-elt-2* as the 2E cells gastrulate, showing DAPI (M) and smiFISH signal (N). The nuclei of Ea and Ep are outlined. (O,P) Later expression of *Can-elt-2* in the gut primordium. (Q,R) Onset of *Cel-elt-2* at the 4E stage. The nuclei of the E granddaughters are outlined. (S,T) Later expression of *Cel-elt-2* in the gut primordium. For each probe set, we examined at least 50-100 total embryos. Embryos are approximately 50 μm long. Anterior is to the left, and dorsal is up.

**Fig. 5.**
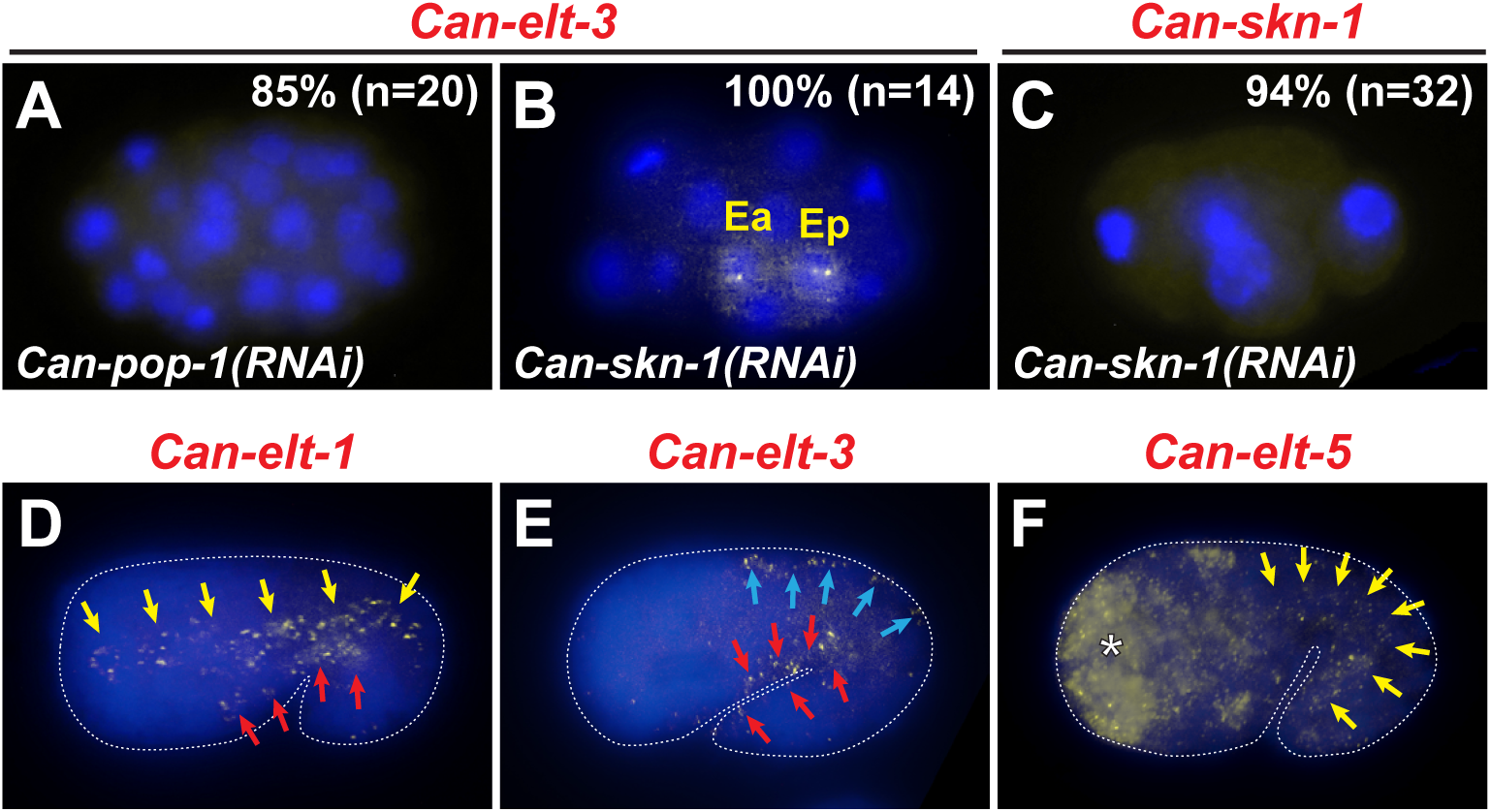
Control smiFISH experiments in *C. angaria*. (A) Depletion of *Can-pop-1* by RNAi results in a loss of expression of *Can-elt-3*. 100% (n=20) of control 8-cell to ∼50-cell embryos showed staining of *elt-3*, while only 15% (n=20) showed staining after *Can-pop-1(RNAi)* by feeding, comparable to the ∼90% RNAi efficacy of phenotype that we observed. (B) *Can-elt-3* transcripts were detected in all *Can-skn-1(RNAi)* embryos of appropriate stage (n=14). (C) *Can-skn-1* mRNA was detected in 97% (n=34) of controls but only 6% (n=32) of *Can-skn-1(RNAi)* embryos, showing that *Can-skn-1(RNAi)* efficiently eliminated *Can-skn-1* mRNA. (D-F) Hypodermal expression of *Can-elt-1, Can-elt-3*, and *Can-elt-5* resembles previously reported expression patterns for the orthologous *C. elegans* genes. (D) Expression of *Can-elt-1* in lateral (yellow arrows) and ventral (red arrows) hypodermal cells, similar to prior reports (Page et al., 1997). (E) Later expression of *Can-elt-3* in dorsal (red arrows) and ventral (yellow arrows) hypodermis consistent with conservation of this component of *elt-3* expression with *C. elegans* (Gilleard et al., 1999). (F) *Can-elt-5* expression in lateral seam cells (yellow arrows) and in many cells in the anterior, likely descendants of AB and MS (Koh and Rothman, 2001).

In *C. elegans, elt-1, elt-3*, and *elt-5* expression patterns are dynamic and occur in multiple cell types in the early embryo, but later become restricted to primarily hypodermal lineages (Page et al., 1997; Gilleard and McGhee, 2001; Koh and Rothman, 2001; Hashimshony et al., 2012; Tintori et al., 2016). By smiFISH, *Can-elt-1* and *Can-elt-5* showed expression patterns that resembled those of their *C. elegans* orthologues, suggesting these have been evolutionarily conserved (Fig. 5D,F and data not shown). More importantly, neither *Can-elt-1* nor *Can-elt-5* showed early lineage-specific expression in E.

In contrast to the other GATA factors, transcripts for *Can-elt-3* were detected in the E cell, just after its birth, and the early E descendants up to the 8E stage (Fig. 4E-H). Transcripts were primarily cytoplasmic in most cells, however we regularly saw 1-2 foci of nuclear staining in E, as well as Ea and Ep, likely representing nascent bursts of transcription on the *Can-elt-3* gene itself (Seydoux and Fire, 1994). (The 1-2 foci are consistent with the expected 1:1 ratio of XX to XO embryos, because *C. angaria* is a male-female species.) This suggests that *Can-elt-3* is zygotically activated in the very early E lineage, showing a pattern of expression that overlaps both *Cel-end-3* and *Cel-end-1* in *C. elegans* (Fig. 4I-L). Consistent with conservation of the later ectodermal component of *elt-3* expression, we observed *Can-elt-3* transcripts in putative hypodermal lineages in older *C. angaria* embryos (Fig. 5E). We further observed very low levels of *Can-elt-3* transcripts in 2-cell and 4-cell stage embryos (Fig. 4C and data not shown), at much lower apparent levels than *Cel-skn-1* and *Can-skn-1* (Fig. 4A,B), and which is not seen for *Cel-elt-3* (Fig. 4D). Low levels of *Can-elt-3* mRNAs were detected independently in pre-8-cell stage embryos by RNA-Seq, but at levels ∼10x lower than that of *Can-skn-1* or *Can-pop-1* (Macchietto et al., 2017).

### The putative ‘b’ isoform of Can-elt-3 is expressed only in the endoderm

The annotated structure of *Cel-elt-3* in WormBase (version WS284) includes two major isoforms: A shorter ‘a’ isoform of length 226 amino acids (aa), and a longer ‘b’ isoform of 317 aa that is extended at the amino end (Fig. 1F). A smaller isoform ‘c’ is 18 aa smaller than isoform ‘a’ at its amino end. The ‘a’ and ‘b’ isoforms have been confirmed to be expressed in *C. elegans* by direct mRNA sequencing (Li et al., 2020; Roach et al., 2020). Since the probe sets we used above can detect the common carboxyl end of all putative *Can-elt-3* isoforms, we repeated the smiFISH staining of *Can-elt-3* using probes derived from only the upstream coding sequences specific to the longer isoform. With this ‘b’-specific probe set, we found that the early E lineage component of *Can-elt-3* expression was still observed, however, neither the early maternal transcripts, nor the later hypodermal component of expression, were detected (n>60 embryos, data not shown). Since our earlier probe set detects both the ‘a’ and ‘c’ isoform, we were unable to distinguish between them. These results nonetheless provide evidence that the longer ‘b’ isoform is endoderm-specific.

### Expression of Can-elt-3 in Can-pop-1(RNAi) and Can-skn-1(RNAi) embryos

Since *Can-elt-3* is expressed in the early E lineage and *Can-pop-1(RNAi)* results in the loss of gut, we wished to determine whether *Can-pop-1* acts upstream or downstream of *Can-elt-3* expression. We used smiFISH to detect *Can-elt-3* transcripts in control and *Can-pop-1(RNAi)* treated embryos. We observed *Can-elt-3* expression in the early E lineage in control embryos from the 8-cell to ∼50-cell stage (100%, n=20), but only in a small fraction of similarly staged embryos in *Can-pop-1(RNAi)* (15%, n=20) (Fig. 5A). The small fraction that did show staining is consistent with our prior measurement of ∼10% of embryos that are unaffected by *Can-pop-1(RNA)*. In parallel experiments, we confirmed that knockdown of *Can-skn-1* did not affect *Can-elt-3* expression (14/14 embryos), although it did result in depletion of *Can-skn-1* transcripts in a majority of embryos (19/20) (Fig. 5B,C). We conclude that *Can-pop-1* is required for *Can-elt-3* expression and therefore acts upstream of gut specification, similar to the *pop-1* orthologues in *C. elegans* and *C. briggsae* (Shetty et al., 2005; Lin et al., 2009; Zhao et al., 2010). Our results also suggest that *Can-skn-1* is dispensable for both *Can-elt-3* expression and gut specification in *C. angaria*.

### RNAi of Can-elt-3 frequently results in the complete loss of endoderm

Because *Can-pop-1(RNAi)* results in a penetrant loss of gut, and of early E lineage *Can-elt-3* expression, we hypothesized that knockdown of *Can-elt-3* would result in the loss of gut specification. We attempted RNAi by feeding to target *Can-elt-3* in *C. angaria* but observed >95% normal embryos or larvae, and all had gut (n>200; Fig. 6A-D). However, RNAi by direct gonadal injection of *Can-elt-3* dsRNA into strain PS1010 resulted in a significant number of arrested embryos and larvae in 76/122 (62%) of progeny in a time window 24-48 h after injection. We examined these by DIC microscopy for the presence of birefringent gut granules, ‘fried-egg’ nuclei typical of gut cells, an intestinal lumen, and basement membrane surrounding the intestine. In almost all cases, these features of differentiated gut were completely absent. In a small number of embryos, we observed rare gut-like nuclei, however we could not see a polarized epithelium and gut granule birefringence, and no lumen was visible (Fig. 6H-J). Except for these few cases, arrested embryos and larvae were strongly reminiscent of *C. elegans end-1(ok558) end-3(ok1448)* double null mutants (Fig. 6M-P) (Owraghi et al., 2010). Unlike *Cel-end-1,3(-)*, however, which show variable elongation from ∼2x-3x, arrested *Can-elt-3(RNAi)* embryos tended to be fully elongated. As well, *Cel-end-1,3(-)* embryos often contain internal hypodermis-lined cavities that result from the transformation of E to a C-like cell when *Cel-end-1* and *Cel-end-3* are absent (Sulston et al., 1983; Zhu et al., 1997; Maduro et al., 2005a). Such cavities were not visible in *Can-elt-3(RNAi)*. We repeated the injections into strain RGD1 with similar results (data not shown). The occurrence of a significant number of progeny embryos lacking recognizable gut tissue in *Can-elt-3(RNAi)*, and the superficial resemblance of the phenotype to *C. elegans end-1 end-3* double mutants, provide compelling evidence that *Can-elt-3* is required for gut specification in *C. angaria*.

**Fig. 6.**
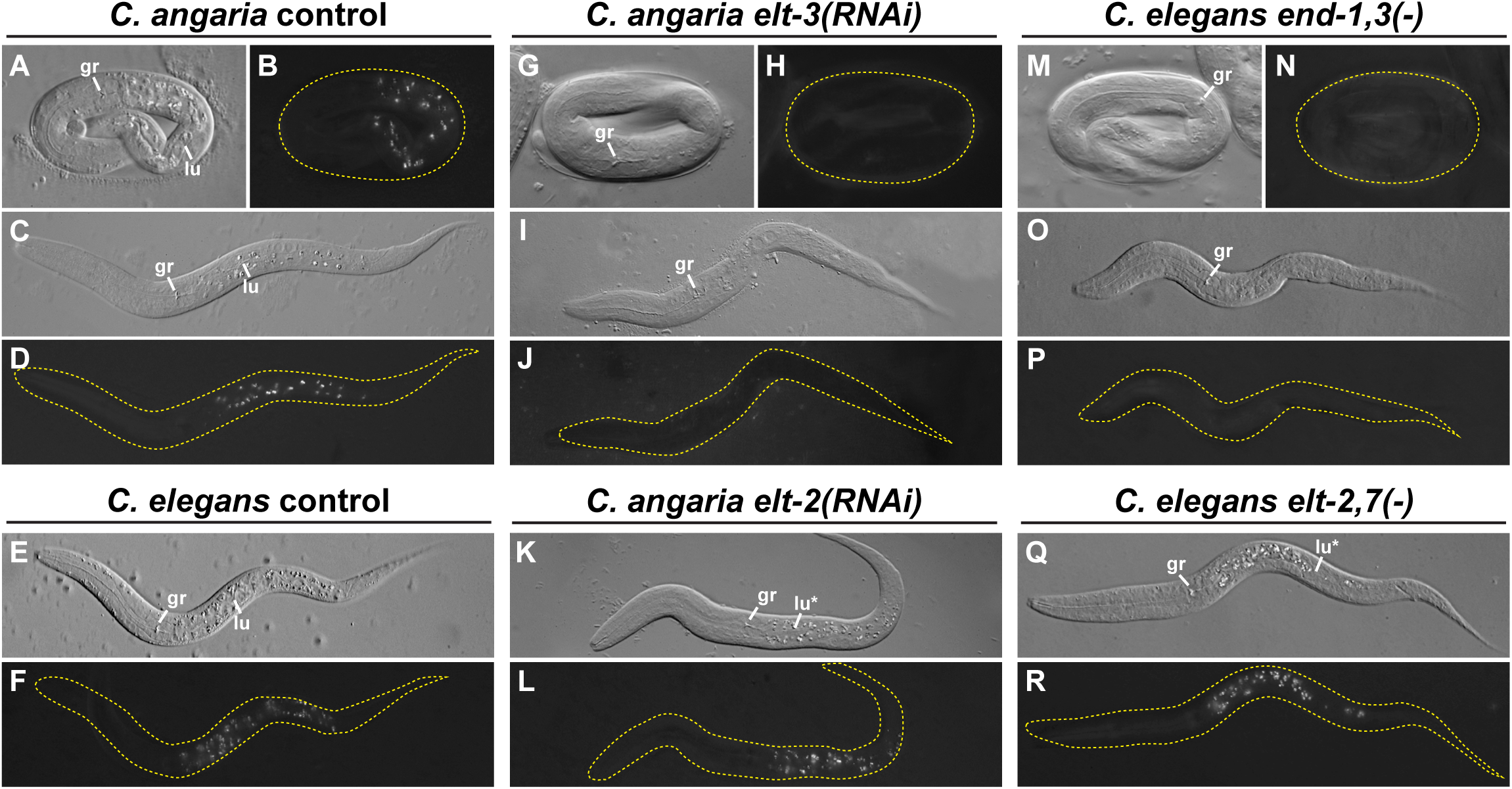
Similarity of specification and differentiation phenotypes between *C. angaria* and *C. elegans*. Images show DIC (gray images) and polarized light (dark images) of representative arrested embryos and larvae following RNA interference of *Can-elt-3* and *Can-elt-2* in *C. angaria*, compared with mutants for *Cel-end-1(ok558) end-3(ok144)* and *Cel-elt-2(ca15); elt-7(tm840)* in *C. elegans*. (A,B) Control *C. angaria* 3-fold embryo. (C,D) Newly hatched *C. angaria* larva. (E,F) Wild-type *C. elegans* newly hatched larva. (G,H) *C. angaria Can-elt-3(RNAi)* 3-fold embryo. (I,J) Arrested *Can-elt-3(RNAi)* showing absence of gut. (K,L) Arrested *Can-elt-2(RNAi)* larva with gut granules but incomplete lumen. (M,N) Arrested 3-fold *Cel-end-1,3(-)* double mutant. (O,P) Absence of gut in arrested *Cel-end-1,3(-)* larva. (Q,R) *C. elegans elt-2(ca15); elt-7(tm840)* arrested larva with abnormal lumen. Abbreviations: gr, grinder in pharynx terminal bulb; lu, example of normal gut lumen; lu*, abnormal or patchy gut lumen. Embryos are approximately 50 μm long. The larvae are shown to the same scale; the control *C. elegans* larva is approximately 240 μm long. In polarized light images, the embryo or larva are outlined with a dashed yellow line. In most images, anterior is to the left and dorsal is up.

### RNAi of Can-elt-2 results in incomplete gut differentiation

We showed earlier that *Can-ELT-2::GFP* can fully complement the larval lethal defects of a *Cel-elt-2; elt-7* double mutant. We hypothesized that *Can-elt-2* has the same function in gut differentiation in *C. angaria* as *Cel-elt-2,7* in *C. elegans*. We therefore examined the effects of *Can-elt-2(RNAi)* using direct dsRNA injection. We observed a penetrant larval lethality in 39/89 (44%) of progeny embryos examined 24-72 h after injection. In these arrested larvae, while intestine was present, we observed a variety of differentiation defects, including a partial intestinal lumen and patches of intestine lacking gut granules (Fig. 6K,L). The phenotype was highly reminiscent of the *C. elegans Cel-elt-2; elt-7* double mutant phenotype (Fig. 6Q,R). We conclude that *Can-elt-2* is required for gut differentiation in *C. angaria*, as expected.

### Overexpression of Can-ELT-3 is sufficient to activate Can-ELT-2::GFP and gut specification in C. elegans

While *Can-elt-3* is necessary to specify gut in *C. angaria*, we wished to test whether expression of *Can-ELT-3* is sufficient to do so. As a proxy for doing this experiment in *C. angaria*, we used *C. elegans*. Prior studies have shown that any of the endodermal GATA factors is able to promote widespread gut specification when they are individually overexpressed (Fukushige et al., 1998; Zhu et al., 1998; Maduro et al., 2001; Maduro et al., 2005a; Sommermann et al., 2010). Because *Can-elt-2* can replace *Cel-elt-2*, we hypothesized that an ability of *Can-ELT-3* to activate *Can-elt-2* could be demonstrated in *C. elegans*. We constructed a heat shock *hs-Can-ELT-3B::CFP* transgene to be able to conditionally express the endoderm-specific ‘b’ isoform throughout early embryos. The transgene was introduced into the *C. elegans elt-2(ca15); elt-7(tm840); Ex[Can-ELT-2::GFP]* strain. We heat shocked mixed-stage early embryos (<100 cells) for 20 min at 34°C. Within 75-90 min, we observed widespread nuclear CFP in

∼60% of embryos, confirming expression of the transgene. After a further 90 min, we began to observe both ectopic onset of *Can-ELT-2::GFP* in ∼40% of embryos, along with diminution (and subsequent absence) of the *Can-ELT-3B::CFP* signal. At the end of embryogenesis, we observed many one-fold arrested embryos containing ectopic gut granules, lacking CFP signal and which showed widespread *Can-ELT-2::GFP* (Fig. 7A-C). We repeated these experiments several times totaling >150 embryos with similar results. In separate experiments, we tested whether the *hs-Can-ELT-3B::CFP* alone, in a wild-type background and the absence of the *Can-ELT-2::GFP* transgene, can promote gut differentiation through endogenous *Cel-elt-2,7*. Consistent with this ability, we observed a high frequency (53%, n=77) of onefold arrested embryos filled with gut-like cells and ectopic gut granules (Fig. 7D,E).

**Fig. 7.**
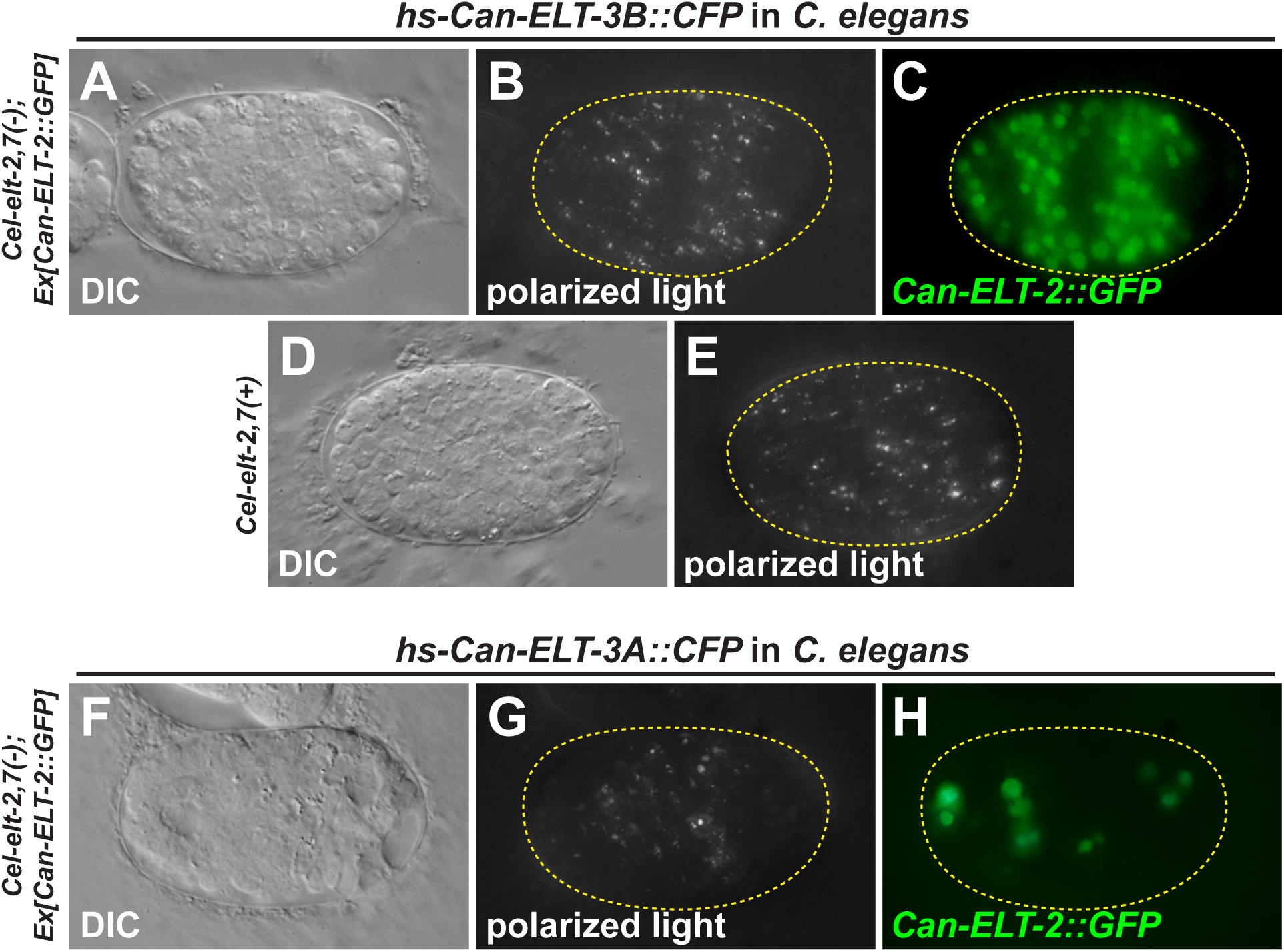
Ectopic overexpression of the *Can-ELT-3* ‘b’ isoform in *C. elegans* activates the *Can-ELT-2::GFP* transgene and promotes gut specification. (A) DIC of a terminal-stage embryo in which *Can-ELT-3B::CFP* was overexpressed by a 20 min heat shock at 34°C of a *hs-Can-ELT-3B::CFP* transgene before the 100-cell stage. (B) Widespread distribution of birefringent gut granules of the same embryo. (C) Widespread expression of the *Can-ELT-2::GFP* transgene in the arrested embryo. (D) DIC image of *hs-Can-ELT-3B::CFP* embryo in a wild-type background. (E) Gut granules in the same embryo in (D). (F) DIC of terminal-stage embryo following *Can-ELT-3A::CFP* overexpression. (G) Apparent granule-like material by polarized light. (H) *Can-ELT-2::GFP* expression of the same embryo in (F) and (G). The location of expression does not match the location of the granules.

We also tested whether the shorter isoforms of *Can-ELT-3* could promote ectopic gut specification. Both *hs-Can-ELT-3A::CFP* and *hs-Can-ELT-3C::CFP* showed induction following heat shock as evidenced by widespread nuclear CFP, and resulting in embryonic arrest in both cases. With the shortest ‘c’ isoform, we failed to observe ectopic expression of *Can-ELT-2::GFP*, and did not see any evidence of ectopic specification of gut at the end of development (n>50 embryos; data not shown). With the ‘a’ isoform, we also did not observe ectopic induction of *Can-ELT-2::GFP* at the expected time compared with overexpression of the ‘b’ isoform. Most embryos arrested with either no gut or a small patch of gut (n=85%, n=39), and a small number showed dispersed gut granules and *Can-ELT-2::GFP*-expressing nuclei (15%, n=39; Fig. 7F-H). Taken together, these experiments show that overexpression of the *Can-ELT-3* ‘b’ isoform in *C. elegans* can efficiently promote gut specification, through either the *Can-ELT-2::GFP* transgene, or endogenous *Cel-elt-2,7*.

Given the ability of *Can-elt-3* to activate *Can-elt-2* in *C. elegans* outside of the gut lineage, one can predict that expression of *Can-elt-3* within the early E lineage alone, driven by one of the *end* promoters, would restore gut specification to a strain lacking *Cel-elt-7* and *Cel-end-1,3* as in the prior results with forced early *Cel-elt-2* and *Cel-elt-7* (Wiesenfahrt et al., 2015; Dineen et al., 2018). Our preliminary experiments along these lines show that this is in fact possible, and a report on these results will be presented elsewhere (G.B.-M. and M.M., unpublished results).

### Use of ELT-3 to specify gut may be widely conserved

Finally, we wished to determine whether other *Caenorhabditis* species outside the Elegans Supergroup might also use *elt-3* to specify gut. We examined expression of *elt-3* and *elt-2* in *C. portoensis*. As shown in Fig. 8A,B, we detected *Cpo-elt-3* and *Cpo-elt-2* in *C. portoensis* embryos in the early E lineage and later gut, respectively. Hence, in at least one other species outside of the Elegans Supergroup, ELT-3 likely has a conserved role in gut specification.

**Fig. 8.**
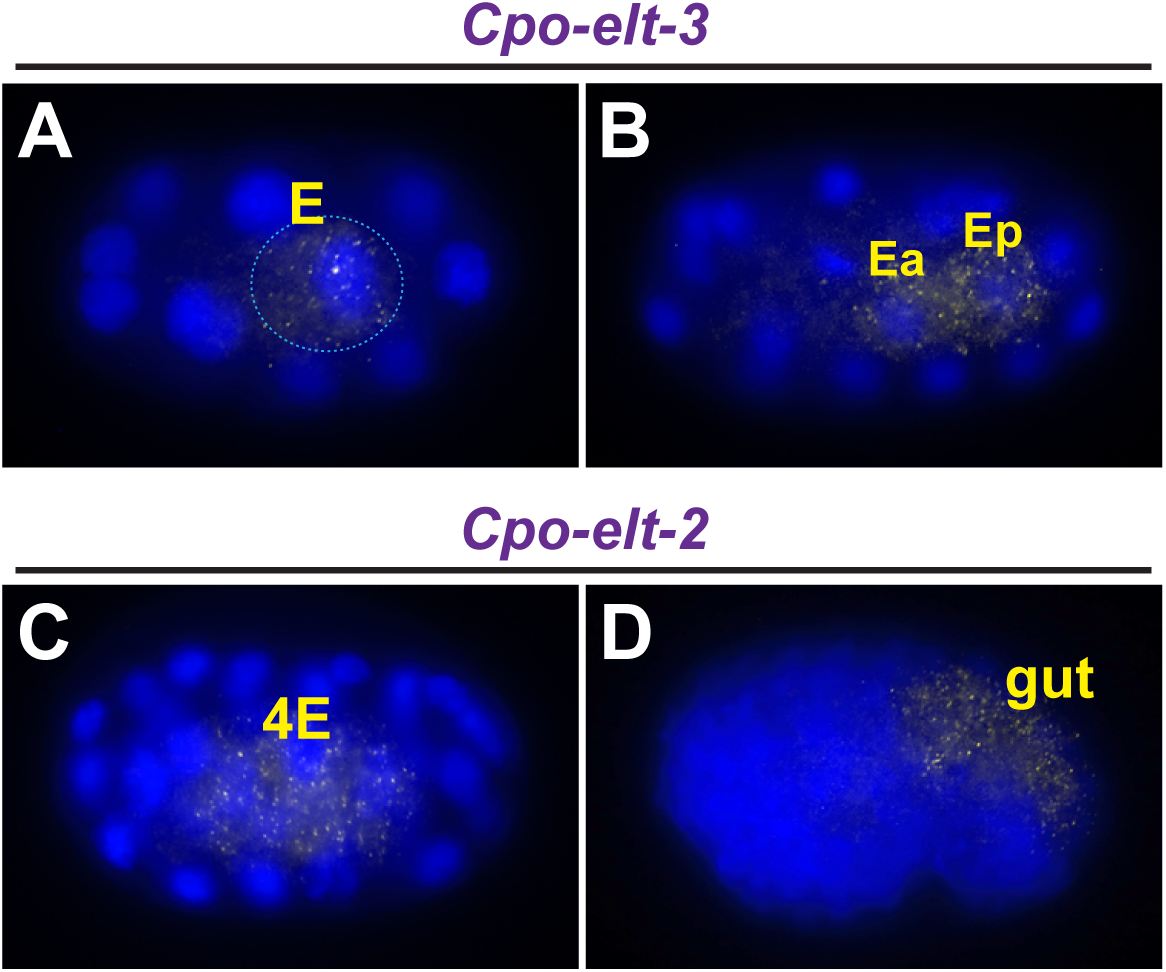
Expression of *elt-3* and *elt-2* orthologues in *C. portoensis* by smiFISH. (A) *Cpo-elt-3* transcripts in E in a 12-cell embryo. (B) *Cpo-elt-3* at 2E stage. (C) *Cpo-elt-2* at 4E stage. (D) *Cpo-elt-2* in bean stage. A *C. portoensis* embryo is approximately 50 μm long.

## Discussion

In this work we have elucidated the core components of an ancestral mechanism for gut specification in *Caenorhabditis*. We began by showing that a *Can-ELT-2::GFP* transgene is expressed in the early E lineage downstream of END-1,3, and is capable of replacing endogenous *Cel-elt-2,7*. We have established that *Can-elt-3* fulfills the criteria of being the *Can-elt-2* activator in *C. angaria*, as it is expressed at the correct time and place; it is necessary for specification of gut; and it is sufficient for specification of gut. Within *C. angaria*, the ‘b’ isoform of *Can-ELT-3* is expressed upstream of *Can-elt-2* in the early E lineage, like the early E lineage expression of *Cel-end-1,3 C. elegans*. Depletion of either *Can-pop-1* or *Can-elt-3* by RNA interference in *C. angaria* results in a penetrant loss of gut specification, with *Can-pop-1* acting upstream of *Can-elt-3* expression. And finally, overexpression of *Can-ELT-3B* throughout *C. elegans* can induce both expression of *Can-ELT-2::GFP* and differentiation of gut. Our results suggest that the ancestral state of the endoderm specification network, as represented by *C. angaria*, consisted of a simpler cascade of maternal POP-1 activating expression of *Can-elt-3*, followed by *Can-ELT-3* activating *Can-elt-2* expression. Differentiation is then activated by *Can-ELT-2* which also maintains its expression by positive by autoregulation.

### Origin of the gut network at the base of the Elegans Supergroup

The simpler ancestral network in *C. angaria*, and the derived networks in *C. elegans* and *C. briggsae*, are shown in Fig. 9. What was a simpler network involving only one GATA factor (ELT-3) had to expand, in a very short evolutionary time, to several regulators (ENDs, ELT-7, MEDs), and along the way, the original ELT-3 also had to lose its endoderm specification role. Such a drastic rapid rewiring of the gene network is a bit unexpected, but the structural similarities among the derived factors point to a common origin (Maduro, 2020). We previously suggested that the *end* and *elt-7* genes might have originated from a partial duplication of *elt-2*, through a successive cascade of upstream duplication and intercalation into the network, with the *med*s originating from a duplication of *end-3* (Maduro, 2020). The findings here suggest the more plausible hypothesis that the *end*s and *elt-7* originated as a duplication of *elt-3* and not *elt-2*. Of the *C. elegans* GATA factors, ELT-3 has features that make it more “endodermal” than hypodermal. For one, it has an intron in the same position as the endodermal GATA factors (*med-1, med-2, end-1, end-3, elt-2, elt-7*), splitting the asparagine codon (N) immediately after the third cysteine of the zinc finger (Maduro, 2020). The *elt-1* and *elt-5* genes lack this intron and instead have one farther downstream in the basic domain. Furthermore, the short carboxyl end of ELT-3, which terminates abruptly after the basic domain, is a feature found only among the MED and END factors and ELT-7, as ELT-1, ELT-2, and ELT-5 contain extended regions after the basic domain (Maduro, 2020).

**Fig. 9.**
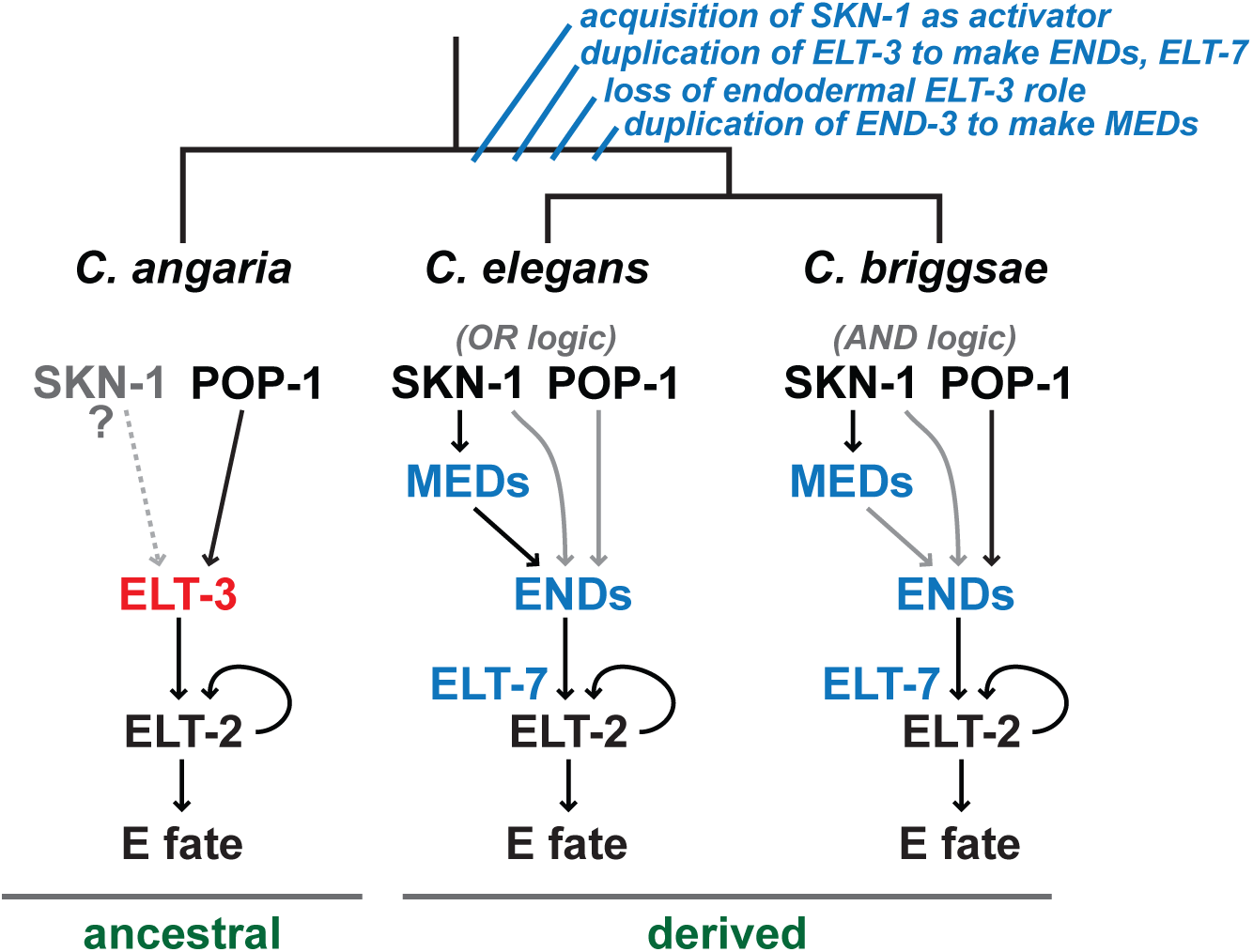
Developmental system drift in endoderm specification in *Caenorhabditis*. Core endoderm specification pathways among *C. angaria, C. elegans*, and *C. briggsae*. In an apparently short evolutionary time, gene duplications followed by divergence of promoter and protein sequences occurred at the base of the Elegans supergroup: ELT-3 likely gave rise to the END factors and ELT-7, then duplication of END-3 gave rise to the MED factors. At some point, the SKN-1 factor becomes a main activator of gut specification, in part through the MED factors, in parallel with POP-1. Arrows show positive regulation, with black arrows indicating stronger inputs, while gray arrows indicate weaker inputs.

### Change in function of maternal regulators compared to C. elegans

POP-1/TCF is a regulator of fates in multiple places in *C. elegans*, where sister cells, born along the anterior-posterior axis (including MS and E), adopt different fates due to the state of POP-1 (Lin et al., 1995; Lin et al., 1998). In anterior cells like MS, POP-1 acts as a repressor, while in posterior cells, POP-1 nuclear concentrations are reduced, and it is adopts an ‘activator’ state through interaction with the divergent β-catenin, SYS-1 (Huang et al., 2007; Phillips et al., 2007). In the case of the MS-E fate decision, the activator state in E is the result of an extrinsic Wnt/MAPK/Src signal that results from contact between EMS, the mother of MS and E, and the neighboring cell, P_2_ (Goldstein, 1992; Goldstein, 1993; Rocheleau et al., 1997; Thorpe et al., 1997; Bei et al., 2002). In *C. elegans*, loss of maternal *Cel-pop-1* results in a phenotype in which MS adopts an E-like fate and produces gut (Lin et al., 1995). A positive contribution of POP-1 to E specification is detectable only when other regulatory inputs, such as *Cel-skn-1* or *Cel-end-3*, or the *Cel-med*s, are also eliminated, in which case loss of *Cel-pop-1* will enhance the loss-of-gut phenotype (Maduro et al., 2005b; Maduro et al., 2007). We have indicated this in Fig. 9 and a prior work as ‘OR logic’ (Lin et al., 2009). POP-1 as an effector of Wnt signaling generally acts in this manner in other contexts, making the phenotype of loss of *Cel-pop-1* in gut specification non-canonical (Rocheleau et al., 1997; Thorpe et al., 1997; Rocheleau et al., 1999; Thorpe et al., 2000; Lam and Phillips, 2017). By contrast, in *C. briggsae*, depletion of *Cbri-pop-1* by RNAi has a canonical Wnt phenotype and results in the penetrant loss of gut, just as with the *Can-pop-1(RNAi)* phenotype we observed here (Lin et al., 2009; Zhao et al., 2010; Nusse and Clevers, 2017). This is indicated as ‘AND logic’ in Fig. 9 (Lin et al., 2009). Hence, it is conceivable that the elimination of gut when *pop-1* is depleted is the ancestral state, retained in *C. briggsae*, but derived in *C. elegans*. This derived state may not be the result of a change in POP-1 *per se*, but rather the dynamic changes that occur within the downstream network within the Elegans Supergroup. For example, the change in POP-1 state between *C. elegans* and *C. briggsae* may be the result of an apparent increased role for the MEDs in activation of *Cel-end-3*, as the *Cbri-end-3* orthologues have fewer MED sites in their promoters, and indeed *C. elegans* has four such sites, which is high among the Elegans Supergroup species (Broitman-Maduro et al., 2005; Maduro, 2020).

In contrast with POP-1, the loss of *Can-skn-1* by RNAi failed to produce an effect. We controlled for the effectiveness of the RNAi treatment by examining embryos for loss of *Can-skn-1* mRNA, suggesting that the failure to see a phenotype is either the result of a loss of a role for *Can-SKN-1* in E and MS fates, or another parallel regulator that provides redundant function (indicated by a ‘?’ in Fig. 9). We looked for, but did not find SKN-1 binding site clusters in the *Can-elt-3* upstream region, of the type seen in *med* genes within the Elegans Supergroup (data not shown) (Maduro et al., 2001; Maduro, 2020). Such sites might result in precocious expression of *Can-elt-3* in the late EMS cell, similar to *Cel-med-1,2*, rather than in the E cell as we observed here (Maduro et al., 2007; Raj et al., 2010). Even among *C. elegans* wild isolates, loss of *skn-1* varies in its phenotypic effect on gut specification, from a strong effect to an almost negligible effect, indicating a cryptic effect of polymorphisms distributed throughout the genome (Torres Cleuren et al., 2019). In *C. briggsae*, loss of *Cbri-skn-1* by RNAi resulted in a stronger endoderm phenotype (∼100% loss of gut) than what is seen in the laboratory *C. elegans* strain N2 (∼80% loss of gut) (Bowerman et al., 1992; Lin et al., 2009). Hence, SKN-1 function in the early embryo appears to evolve rapidly within the Elegans Supergroup. An important difference between species in the Elegans Supergroup and *C. angaria*, however, is the absence of the MED factors, which act as zygotic effectors of SKN-1 function, and are actually more important for specification of MS rather than E (Maduro et al., 2001; Maduro et al., 2007). Since all Elegans Supergroup species contain *med* genes, it may be that the recruitment of SKN-1 into MS and E specification requires *med*s.

The ancestral role of SKN-1 is not likely to be in the endoderm. SKN-1 is known to be a major effector of postembryonic responses to physiological stress, and in an unexpected convergence of function, *Cel-ELT-3* interacts genetically with *Cel-SKN-1* in coregulation of genes in oxidative stress responses (Blackwell et al., 2015; Hu et al., 2017). As SKN-1/Nrf proteins are known to be widely conserved in stress responses, it is likely this function of SKN-1 that is ancestral (Blackwell et al., 2015). It may be, then, that the role of SKN-1 in embryonic cell specification is derived in the Elegans Supergroup.

Given that SKN-1 and POP-1 have roles in other contexts, and show dynamic changes in their roles in gut specification even within the genus, we expect that even among nematode species outside the Elegans supergroup, there are likely to be species-specific differences in their roles resulting from developmental system drift (True and Haag, 2001; Lin et al., 2009; Ewe et al., 2020).

### Fates of E in Can-pop-1(RNAi) and Can-elt-3(RNAi)

In the species in which it has been studied, failure of specification typically results in ‘transformed’ blastomeres in which the cells adopt a coherent fate like one of the other early embryonic cells. For example, in *C. briggsae, Cbri-pop-1(RNAi)* results in E adopting an MS-like fate, producing muscle and pharyngeal cells (Sulston et al., 1983; Lin et al., 2009; Zhao et al., 2010). In *C. elegans*, loss of *end-1* and *end-3* together results in a transformation of E to a C-like cell, producing muscle and hypodermis (Zhu et al., 1997; Maduro et al., 2005a). We looked for evidence of ectopically produced tissues that might result from the descendants of an E lineage that does not produce endoderm in *Can-pop-1(RNAi)* and *Can-elt-3(RNAi)* embryos but have not yet found anything conclusive. In *C. elegans*, ectopic cells made by E in gut specification mutants frequently migrate to where these tissues are normally made, which can make it difficult to find them in intact arrested embryos (Choi et al., 2017). In the case of an *end-1 end-3* double mutant, we have occasionally seen internal cavities lined with apparent hypodermis, but these were not seen in either *Can-pop-1(RNAi)* or *Can-elt-3(RNAi)* embryos (Owraghi et al., 2010). Cell fate transformations, at least in *C. elegans*, arise from the co-presence of factors that can specify different blastomere identities, but have a hierarchy. For example, SKN-1 and PAL-1, a specifier of C fate, are both found in the EMS cell, the mother of MS and E (Sulston et al., 1983; Bowerman et al., 1993; Hunter and Kenyon, 1996). When *skn-1* function is absent, PAL-1 is now able to promote a C-like fate from both MS and E (Hunter and Kenyon, 1996). In *C. angaria*, we were unable to find a function for *Can-skn-1*, hence it is not clear if a ‘wholesale’ transformation of fate will necessarily occur. Indeed, in a *C. elegans end-1,3(-); pop-1(-)* embryo, the E cell adopts a mixed fate consisting of both MS-like and C-like cells (Owraghi et al., 2010). Even in *C. elegans*, a failed specification need not result in a fate transformation, for example loss of PAL-1 results in the C blastomere adopting no recognizable fate (Hunter and Kenyon, 1996). Alternatively, it may be that we are not seeing a complete fate transformation because of an incomplete RNAi effect of *Can-elt-3(RNAi)*. We note that incomplete RNAi efficacy was also observed following dsRNA injection targeting *end-1* and *end-3* simultaneously in *C. elegans* (Maduro et al., 2005a). We hypothesize that a chromosomal knockout of *Can-elt-3* would have a completely penetrant phenotype. Hence, determining what happens to the transformed E descendants may have to wait for a *bona fide* mutation of *Can-elt-3*, and future studies to define a more comprehensive set of cell fate regulators.

### Differential activity of ELT-3 function encoded upstream of the DNA-binding domain

We have found that in *C. angaria* the *Can-elt-3* gene has very low-level maternal expression, strong expression in the early E lineage, and later expression in the hypodermis. Our smiFISH results show that the longer ‘b’ isoform is expressed specifically in the early E lineage, from which we deduce that it is this isoform that is the predominant E-specifying gene product. Consistent with this hypothesis, we found that overexpression of the longer ‘b’ isoform of *Can-ELT-3* can efficiently activate *Can-ELT-2::GFP* and promote gut fate in *C. elegans* (Fig. 7). These findings reconcile an apparent paradox with a prior report that *Cel-ELT-3* overexpression promotes ectodermal fates (Gilleard and McGhee, 2001). These experiments used only the smaller ‘a’ isoform known at the time (Gilleard et al., 1999). Similarly, forced expression of the *Cel-ELT-3* ‘a’ isoform within the early E lineage was unable to specify gut (Wiesenfahrt et al., 2015). We also failed to find efficient ectopic gut specification with overexpression of the *Can-ELT-3* ‘a’ isoform.

We further hypothesize that the ability of *Can-ELT-3B* to activate *Can-elt-2* and *Cel-elt-2* resides, at least partly, in the amino portion of the protein specific for the long isoform (Fig. 1F). This part of the protein is less conserved with *Cel-ELT-3B*, save for a short 21-23 amino acid region at the anterior terminus that contains a short ‘poly-S/T’ motif of unknown significance (Maduro et al., 2005a; Maduro, 2020). In human and mouse, protein-protein interactions outside of the DNA-binding domains and combinatorial interactions at promoters explain differential activities of GATA factors (Romano and Miccio, 2020). Hence, it is highly plausible that amino-terminal portion of the *Can-ELT-3B* isoform interacts with cofactors or is important for a pioneering role in establishing an active transcription state of *Can-elt-2*. A role for GATA3 as such a pioneering factor was recently described (Tanaka et al., 2020).

### Remnant expression of Cel-elt-3 in the C. elegans endoderm

*Cel-ELT-3* was initially associated with only the hypodermis (Gilleard et al., 1999). Our observation of continued hypodermal expression of *Can-elt-3* suggests this component of expression/function for ELT-3 is both ancestral and conserved. However, a null mutation of the gene has no detectable phenotype, showing that its hypodermal function is non-essential (Gilleard and McGhee, 2001). There is some evidence for endodermal expression of *Cel-elt-3* both embryonically and post-embryonically. In the early E lineage, though not in E itself, a low level of *Cel-elt-3* transcripts has been observed by single-cell transcriptomics (Hashimshony et al., 2012; Tintori et al., 2016). Given that *elt-7* and *end-1* single mutants also have no apparent phenotype, we looked for a cryptic role for *elt-3* in gut specification by combining a putative null mutant, *elt-3(gk121)*, with each of *end-1(ok558), end-3(ok1448)*, and *elt-7(tm840)*. However, in all cases, no synergistic enhancement of phenotype was seen, suggesting that either no protein is made from these transcripts, or expression is dependent on one or more of these factors (G.B.-M. and M.M., unpublished results). Hence, there does not seem to be any remnant contribution of *Cel-ELT-3* to endoderm specification, suggesting this function has been completely replaced by the MED/END-1,3/ELT-7 factors in the Elegans Supergroup.

Later expression of *Cel-elt-3* in the intestine has been reported, though this has been controversial (Budovskaya et al., 2008; Tonsaker et al., 2012). At least one image from the TransgeneOme project of a *elt-3::TY1::EGFP::3xFLAG* translational reporter shows nuclear intestine expression (Fig. S1) (Sarov et al., 2012). Potentially, intestinal expression of *elt-3* in the later intestine could be conditional, arising in response to oxidative stress as it appears to function in the hypodermis (Budovskaya et al., 2008; Shao et al., 2013; Hu et al., 2017).

### A simpler network for gut specification

An unexpected aspect of the putative ancestral gut specification network is its simplicity. It is reminiscent of gut development in *Drosophila*, in which two GATA factors act in a similar cascade: *Serpent (srp)* specifies gut fate upstream of *dGATAE*, which executes and maintains this fate (Reuter, 1994; Okumura et al., 2005). Within *C. elegans*, then, why bother with five specification factors (MED-1, MED-2, END-1, END-3, ELT-7) when one will do? As hypothesized in a prior work, the expanded network in the Elegans Supergroup may have resulted from evolutionary pressure to accelerate development (Maduro, 2020). Indeed, early developmental timing events are slightly accelerated in *C. elegans* relative to *C. angaria*, although later developmental milestones are similarly timed (Macchietto et al., 2017).

Although paradoxical, an increased number of regulators could amplify early specification and assure rapid, robust activation of *elt-2*, permitting development to speed up without sacrificing robustness (Maduro, 2020). Indeed, we previously compared the expansion of genes in the Elegans Supergroup endoderm network with the emergence of *bicoid* in *Drosophila*, in which expansion of a gene network allowed a more rapid embryonic development to occur (McGregor, 2005). A detailed study of the dynamics of *elt-3* expression between ancestral species like *C. angaria* and species within the Elegans Supergroup would allow this question to be studied.

## Materials and methods

### Identification of orthologous genes in C. angaria

Orthologous GATA factors were identified by BLAST searches as in (Maduro, 2020) using a high-quality genome sequence of PS1010 generously provided by Taisei Kikuchi, University of Miyazaki, Japan. Identity of each GATA factor was confirmed using features identified in a prior work, which include location of introns in the coding region, and signature amino acids within the DNA binding domains (Eurmsirilerd and Maduro, 2020). For other genes we made use of gene predictions from a previously published sequence of *C. angaria* (Macchietto et al., 2017) and from its close relative *C. castelli* (downloaded November, 2020 from caenorhabditis.org). Sequences and gene models are available in Supplemental Data File S1.

### Caenorhabditis strains and transgenesis

The *C. angaria* strains used were PS1010 and RGD1. The *C. portoensis* species was EG4788. *C. elegans* strains were constructed by standard crosses and microinjections to generate transgene arrays (Brenner, 1974; Mello et al., 1991). Mutations: *LG III: unc-119(ed4). LG IV: him-8(e1489). LG V: elt-7(tm840), end-1(ok558), end-3(ok1448). LG X*: *elt-2(ca15)*.

Genotypes were confirmed using allele-specific primers. Transgene arrays: *irEx498 [end-3(+) (pMM768), unc-119::mCherry (pMM824)], irEx798 [Can-elt-2p::ELT-2genomic::GFP::elt-2_3’UTR (pGB598), unc-119::CFP (pMM809), unc-119(+) (pMM016B)], irEx804 [Cel-end-3p::END-3genomic::Can-ELT-3_DNA-binding domain::CFP::Cel-end-3_3’UTR (pGB612), unc-119::YFP (pMM531), unc-119(+) (pMM016B)], irEx807 [hsp16-41::Can-ELT-3(isoform_c)::CFP (pGB614), rol-6D (pRF4)], irEx808 [hsp16-41::Can-ELT-3(isoform_b)::CFP (pGB619), rol-6D (pRF4)], irEx809 [hsp16-41::Can-ELT-3(isoform_a)::CFP (pGB620), rol-6D (pRF4)]*.

### Cloning of transgenes

We constructed transgenes using Gibson assembly (Gibson et al., 2009). The coding region for GFP was amplified from pPD95.67 (a gift from Andrew Fire). Plasmids containing coding regions for the fast-folding fluorescent proteins sCFP3A and Venus/YFP (Balleza et al., 2018) were obtained from Addgene (www.addgene.org). Partial or complete coding regions for *Can-elt-3, Can-skn-1* and *Can-pop-1* were synthesized by IDT (www.idtdna.com).

### Overexpression of Can-ELT-3 by heat shock

We used Gibson assembly to construct an intronless heat shock *Can-ELT-3B::CFP* transgene in vector pPD49.83. We injected the *hs-Can-ELT-3B::CFP* transgene along with the *rol-6*^*D*^ marker (plasmid pRF4) into the *elt-2(ca15); elt-7(tm840)* genetic background rescued with the *Can-ELT-2::GFP* transgene. We constructed a *hs-Can-ELT-3A::CFP* transgene derived from the coding DNA used to construct the ‘b’ isoform transgene, and a smaller heat shock transgene corresponding to the ‘c’ isoform of *Cel-elt-3*, which starts 18 amino acids after the start of isoform ‘a’, using genomic DNA as a template. Both *Can-ELT-3A* and *Can-ELT-3C* were expressed after heat shock as confirmed by widespread nuclear CFP after heat shock, however there was no ectopic activation of *Can-ELT-2::GFP*.

### RNA interference

For RNAi experiments, either genomic DNA fragments or synthesized cDNA sequences were cloned into the feeding-based RNAi vector pPD129.36. We used primers L4440A (gagcgcagcgagtcagtgagcg) and L4440B (cccagtcacgacgttgtaaaacg) to PCR-amplify a template for synthesis of RNA using the T7 MEGAscript kit (ThermoFisher). dsRNA at a concentration of ∼2 μg/μL was injected into one gonad arm per female. To prevent possible cross-interference with mRNA of the other GATA factors, we targeted sequences upstream of the coding regions for the DNA-binding domains. Sequences use for RNAi by feeding or dsRNA injection are in Supplemental Data File S2.

### Microscopy and imaging

Images were obtained using either a Canon EOS 77D or Canon EOS RP camera with an LMscope adapter (Micro Tech Labs, Austria) on an Olympus BX-51 fluorescence microscope equipped with DIC optics. Images were processed for contrast and color using Adobe Photoshop. Fluorescence images were made by combining multiple images taken at different focal planes.

### Detection of RNA in situ

We used the smiFISH protocol (Tsanov et al., 2016; Calvo et al., 2021) adapted for use in *Caenorhabditis* by (Parker et al., 2021) using our previously described fixation protocol (Broitman-Maduro and Maduro, 2011). Probe sets of 16-24 nucleotides plus FLAP-X sequence of (Tsanov et al., 2016). These were generated using the Stellaris Probe designer (www.biosearchtech.com) and are listed in Supplemental Data File S2. Conjugated X FLAP oligos (CACTGAGTCCAGCTCGAAACTTAGGAGG) that were 5′ and 3′ end-labelled with Quasar 570 or Cal Fluor 610 were synthesized by Biosearch Technologies. FLAP-X oligos 5′ and 3′ end labelled with Cy5 or Cy3 were synthesized by IDT. To detect fluorescent smiFISH signals, we used filter sets obtained from Chroma as follows. For Quasar 570, we used the Gold FISH 49304 ET set, and for Cy5, the Narrow-Excitation Cy5 49009 ET set. Rhodamine set 31002 was sufficient for detection of Cal Fluor 610. We imaged co-stained embryos that had both Quasar 570 and Cy5 probes in order of increasing wavelength, i.e. DAPI → Gold FISH → Cy5, to prevent imaging of photoconverted Cy5 (Cho et al., 2021). In our hands, staining was highly consistent among a set of fixed embryos, such that when signal was detected in embryos of a particular stage, signal was seen in all other embryos of that stage. Rare embryos (<5%) that did not show staining were usually visibly damaged and were more likely to be younger than 4-cell stage. A notable exception was detection of *Can-elt-2* transcripts in the *Can-ELT-2::GFP* strain, in which ∼60% of embryos showed staining, consistent with the transmission frequency of the transgenic array. We performed smiFISH following RNAi by feeding in *C. angaria* and included controls for permeabilization and staining in each case. For *Can-pop-1(RNAi)*, we simultaneously stained for *Can-elt-3* using Quasar 570, and *Can-eef1A*.*1*, the orthologue of *Cel-eef1A*.*1* (also known as *Cel-eft-3*), using Cy5. From single-embryo RNA-Seq data, *Can-eef1A*.*1* is expressed at all embryonic stages from zygote through hatching (Macchietto et al., 2017). For *Can-skn-1(RNAi)*, we stained for *Can-skn-1* using Quasar 570 and *Can-elt-3* using Cy5. Because *Can-skn-1(RNAi)* did not result in a loss of gut specification, we reasoned that *Can-elt-3* expression would be unaffected and hence this serves both as confirmation of this hypothesis as well as a control for staining of *Can-skn-1* transcripts following *Can-skn-1(RNAi)*.

## Acknowledgements

We are grateful for pre-publication access to a high-quality PS1010 sequence from Taisei Kikuchi, University of Miyazaki, Japan; and Jordan Ward, UC Santa Cruz for very helpful suggestions. Some strains were provided by the CGC, which is funded by the NIH Office of Research Infrastructure Programs (P40 OD010440).

## Data availability

This work did not produce datasets that need deposition.

